# Chd1 regulates repair of promoter-proximal DNA breaks to sustain hypertranscription in embryonic stem cells

**DOI:** 10.1101/659995

**Authors:** Aydan Bulut-Karslioglu, Hu Jin, Marcela Guzman-Ayala, Andrew JK Williamson, Miroslav Hejna, Anthony D Whetton, Jun S. Song, Miguel Ramalho-Santos

**Affiliations:** Eli and Edythe Broad Center of Regeneration Medicine and Stem Cell Research, Center for Reproductive Sciences and Diabetes Center, University of California, San Francisco, San Francisco, CA 94143, USA; Max Planck Institute for Molecular Genetics, Berlin 14195, Germany; Carl R. Woese Institute for Genomic Biology; Department of Physics, University of Illinois, Urbana-Champaign, Urbana, IL 61801, USA; Department of Biomedical Informatics, Harvard Medical School, Boston, MA 02115, USA; Senti Biosciences, 2 Corporate Drive First Floor, South San Francisco, CA 94080; Stoller Biomarker Discovery Centre, The University of Manchester, Manchester, M20 3LJ, United Kingdom; Thermo Fisher Scientific, Stafford House, HP2 7GE; Lunenfeld-Tanenbaum Research Institute and Department of Molecular Genetics, University of Toronto, Toronto, ON M5G 1X5, Canada

## Abstract

Stem and progenitor cells undergo a global elevation of nascent transcription, or hypertranscription, during key developmental transitions involving rapid cell proliferation. The chromatin remodeler Chd1 binds to genes transcribed by RNA Polymerase (Pol) I and II and is required for hypertranscription in embryonic stem (ES) cells *in vitro* and the early post-implantation epiblast *in vivo*. Biochemically, Chd1 has been shown to facilitate transcription at least in part by removing nucleosomal barriers to elongation, but its mechanism of action in stem cells remains poorly understood. Here we report a novel role for Chd1 in the repair of promoter-proximal endogenous double-stranded DNA breaks (DSBs) in ES cells. An unbiased proteomics approach revealed that Chd1 interacts with several DNA repair factors including Atm, Parp1, Kap1 and Topoisomerase 2β. We show that wild-type ES cells display high levels of phosphorylated H2A.X and Kap1 at chromatin, notably at rDNA in the nucleolus, in a Chd1-dependent manner. Loss of Chd1 leads to an extensive accumulation of DSBs at Chd1-bound Pol II-transcribed genes and rDNA. Genes prone to DNA breaks in Chd1 KO ES cells tend to be longer genes with GC-rich promoters, a more labile nucleosomal structure and roles in chromatin regulation, transcription and signaling. These results reveal a vulnerability of hypertranscribing stem cells to endogenous DNA breaks, with important implications for developmental and cancer biology.

## Introduction

Proliferating stem and progenitor cells are net generators of new cellular biomass and therefore have high biosynthetic demand. One way that which stem/progenitor cells cope with this demand is to enter a state of hypertranscription, which involves a global elevation of nascent transcriptional output^1^. Hypertranscription is masked by most transcriptional profiling approaches, but has attracted renewed interest recently. Hypertranscription has been documented to occur and play critical roles in Embryonic Stem (ES) cells^2^, the post-implantation epiblast^3^, emergence of definitive hematopoietic stem cells^4^, primordial germ cells^5^ and neurogenesis^6^, and may take place in other settings during development, regeneration and disease^1,7^.

The molecular regulation of hypertranscription, or even how it differs from general transcriptional regulation, remains poorly understood. It is expected that hypertranscription involves a coordinated interplay between activating transcription factors, chromatin remodelers and RNA Polymerases. Some of the players implicated in promoting hypertranscription are the transcription factors Myc and Yap/Taz, the RNA Polymerase regulator pTEFb and the chromatin remodeler Chd1 (reviewed in ^1^). Chd1 is an ATP-dependent chromatin remodeler that binds specifically to H3K4me3 and is found at sites of active transcription^8,9^. Chd1 removes nucleosomal barriers to transcriptional elongation^10^ and is required for the optimal activity of RNA Pol I and II^3^. Loss of Chd1 does not affect transcription per se, but it blunts the ability of stem cells to enter hypertranscription in vitro and in vivo^3,4^. Despite these recent insights, the molecular function of Chd1 in hypertranscribing cells remains unclear.

Hypertranscription is a dynamic phenomenon that is responsive to extrinsic cues^2,4,5^, and may therefore share features with ligand-triggered target gene induction. In hormone-responsive cells, target genes are induced via a mechanism that involves the generation of transient endogenous DNA breaks by Topoisomerase II at promoters^11^. Similar induction of target gene transcription mediated by DNA breaks occurs upon exposure to serum or heat shock^12,13^, during zygotic genome activation and in neurogenesis^14,15^. DNA breaks may relieve torsional stress and facilitate DNA unwinding and access of RNA Polymerases^16^. Interestingly, endogenous DNA breaks have recently been shown to occur throughout the genome at the promoters of transcribed genes^17^, suggesting that the link between DNA breaks and transcription may be more general. It remains unknown how cells coordinate the occurrence of DNA breaks and their repair with transcription, a coordination that is anticipated to be of particular importance in hypertranscribing pluripotent cells.

In this study, we report that Chd1 interacts with DNA repair factors in undamaged ES cells. Chd1 promotes the chromatin recruitment/retention of these factors and the repair of DSBs at the promoters of active RNA Pol II-transcribed genes and rDNA in ES cells. Our results reveal an unexpected interplay between Chd1 and the DNA repair associated factors Atm, Kap1 and γH2A.X during the resolution of transcription-associated DSBs in ES cells.

## Results

### Chd1 interacts with double-stranded DNA break repair proteins

In order to probe the function of Chd1 in the regulation of hypertranscription, we identified its interacting proteins by immunoprecipitation followed by mass spectrometry (IP-MS) using a Chd1-Flag knock-in mouse ES cell line (Fig S1A)^3^. We detected 16 of 29 previously described Chd1-interacting proteins that are expressed in ES cells (Fig. 1A). As expected, putative Chd1-interacting proteins are enriched for factors involved in chromatin and transcriptional regulation (Fig. 1B). In addition, there is an unexpected enrichment for DNA repair factors among Chd1 interactors, such as the protein kinase Atm, histone variant H2A.X, MRN complex members Mre11 and Rad50, Parp1/2, Kap1 and topoisomerase 2β (Top2β) (Fig. 1C). We confirmed interactions of Chd1 with activated Atm (Atm phospho-S1981), Kap1, Parp1 and Top2β in wild-type ES cells via co-immunoprecipitation (Co-IP) (Fig. 1D). Interactions of Chd1 with the single-stranded repair factors Xrcc1 and Lig3 were not detected by Co-IP (Fig 1D, data not shown). Therefore, we focused on the double-stranded DNA (DSB) repair components for the remainder of this study.

**Figure 1.**
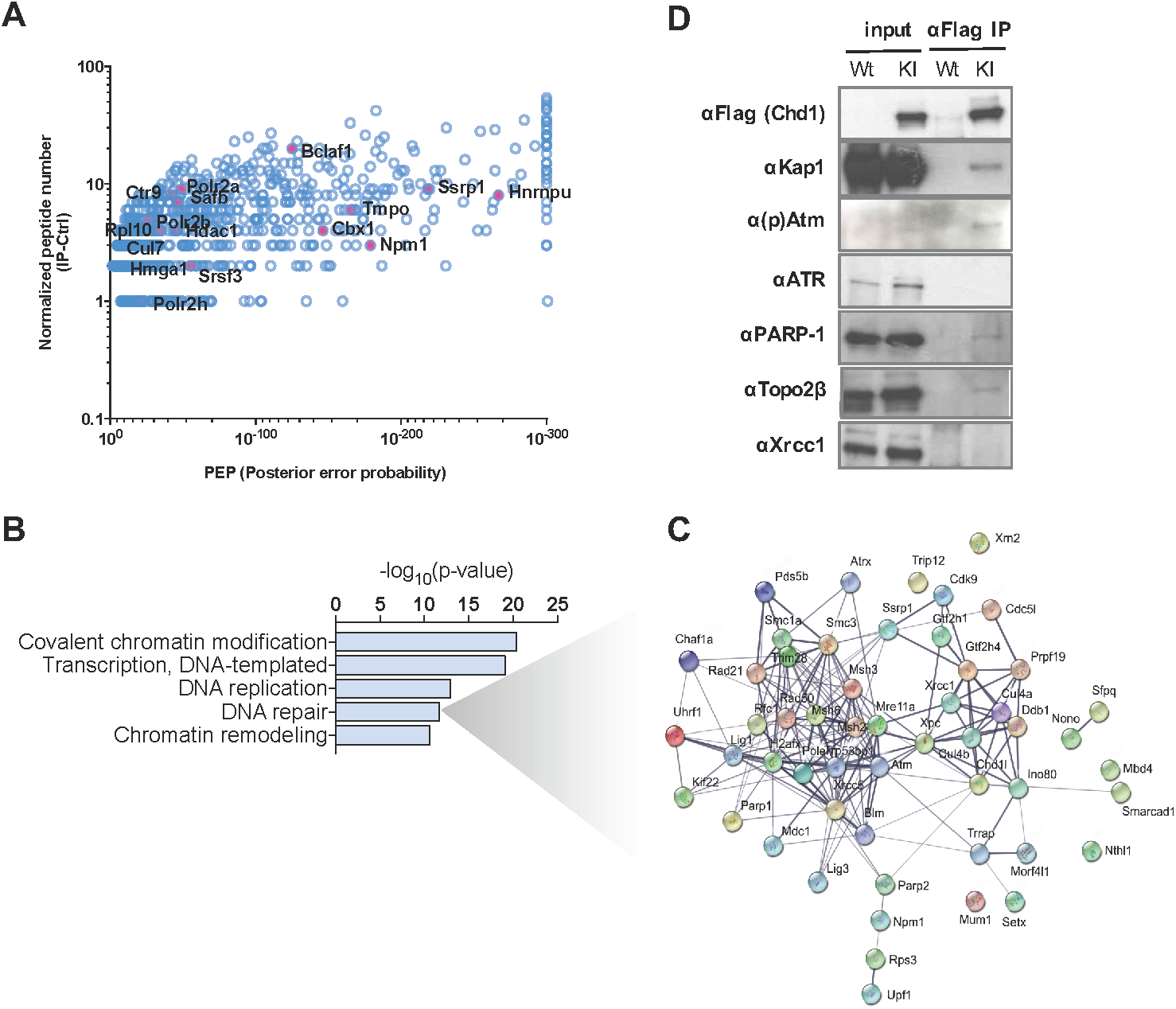
Chd1 interacts with double-stranded DNA repair proteins in ES cells. A) Putative Chd1 interactors, identified by IP-mass spectrometry using a Chd1-Flag knock-in ES cell line. Indicated are previously known interactors of Chd1. B) Gene ontology analysis of co-immunoprecipitated proteins. C) Protein interaction network of factors co-immunoprecipitated with Chd1 and belonging to the gene ontology term “DNA repair”. D) Co-IP validation of the interaction of Chd1 with selected DNA repair proteins.

### DNA repair factors localize to the nucleolus and rDNA in a Chd1-dependent manner

We next assessed how genetic deletion of Chd1^3^ affects the levels and localization of its DSB repair interactors. Loss of Chd1 leads to reduced global levels of S139 phosphorylated H2A.X (γH2A.X), a mark of DSB repair, despite slight increases in the levels of H2A.X and Atm kinase, which phosphorylates H2A.X (Fig. S1B). Immunofluorescence confirmed these results, with the unexpected observation that many of these DNA repair factors accumulate in the nucleolus (Fig. 2A). For example, Kap1 is globally distributed in the nucleoplasm, but phosphorylated Kap1 (pKap1), also a target of Atm with roles in DNA repair^18^, is highly enriched in the nucleolus and reduced in levels in Chd1 KO cells. Atm and Top2β are also primarily localized to the nucleolus, and, in agreement with western blot ting data, their levels increase with Chd1 loss (Fig. 2A). To our knowledge, this pattern of accumulation of DNA repair factors at the nucleolus of undamaged cells has not previously been described. We have previously shown that Chd1 binds directly to rDNA and that loss of Chd1 leads to reduced nascent synthesis of rRNA and fragmentation of nucleoli (^3^ and Fig. 2A). Chromatin immunoprecipitation-qPCR (ChIP-qPCR) confirmed the accumulation of Atm, pKap1 and γH2A.X at rDNA, and the decrease in pKap1 and γH2A.X despite higher levels of Atm in Chd1 KO ES cells (Fig. 2B, S1C). Taken together, these results suggest that Chd1 modulates the function of interacting factors involved in DSB repair in undamaged ES cells, particularly at rDNA in nucleoli.

**Figure 2.**
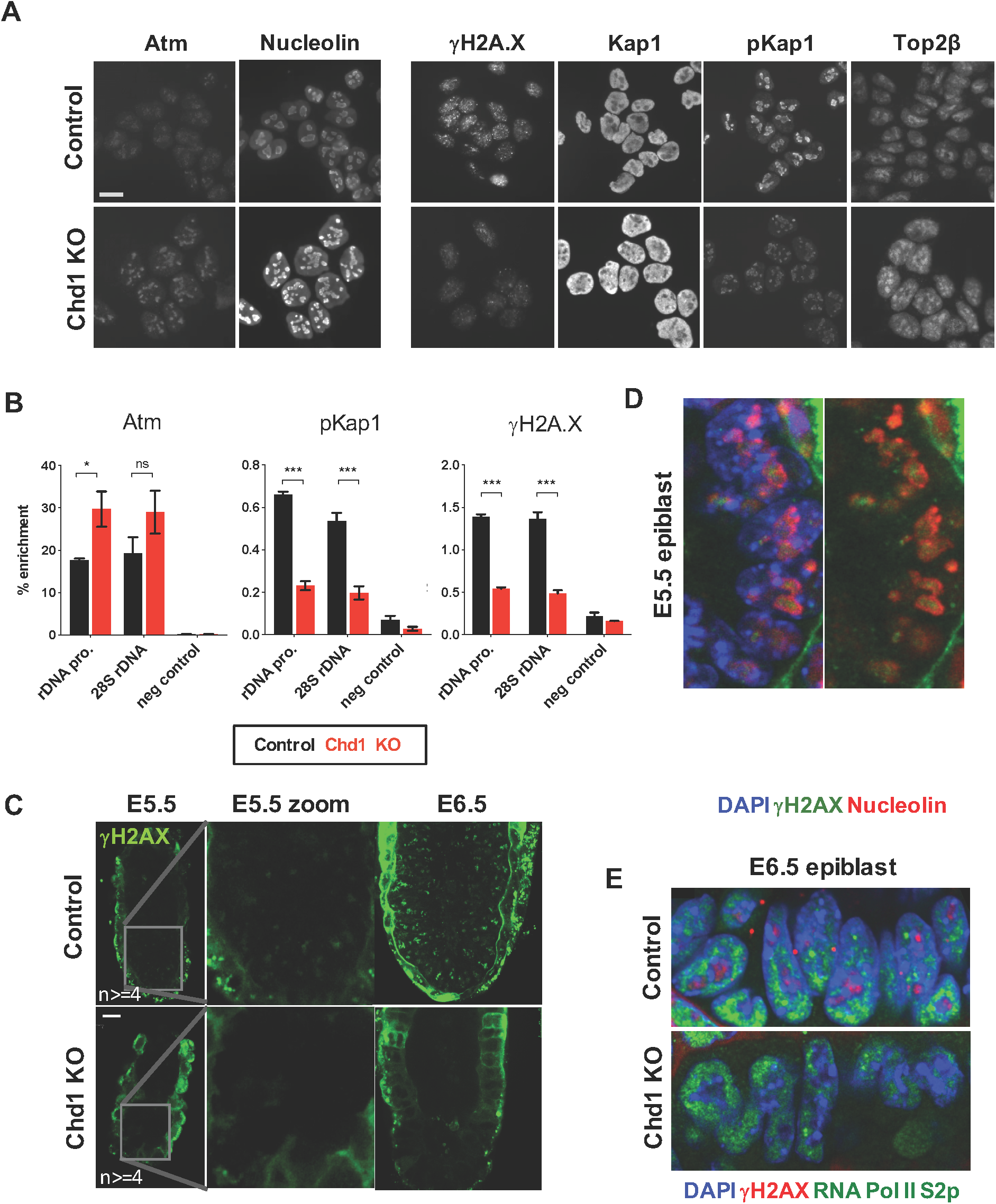
DNA repair signaling is perturbed in Chd1 KO ES cells. A) Immunofluorescence analysis of Chd1-interacting DNA repair proteins in control and Chd1 KO cells. Scale bar denotes 10 μm. Left panel shows Atm-Nucleolin co-staining, right panel shows indicated single stains. B) ChIP-qPCR analysis of selected DNA repair proteins at rDNA. Error bars show standard deviation of two biological replicates. Statistical tests performed are T-tests with Holm-Sidak correction. *, **, *** correspond to p-values of <0.05, 0.01, 0.001. ns= non-significant. C) γH2A.X staining of control and Chd1 KO mouse embryos at E5.5 and E6.5. Scale bar denotes 25 μm. D) γH2A.X and Nucleolin staining of the control E5.5 epiblast, showing nucleolar localization of the γH2A.X signal. E) γH2A.X and elongating RNA Pol II (S2p) staining of the E6.5 epiblast in control and Chd1 KO embryos.

### γH2A.X is present at nucleoli in pre-implantation embryos and is lost in Chd1 KO embryos

We previously used mouse genetics to show that Chd1 is essential for rapid growth of the early post-implantation mouse epiblast (E5.5-6.5) by promoting a high transcriptional output, notably of nascent rRNA at the nucleolus^3^. The surprising presence of γH2A.X in undamaged ES cells and its dependence on Chd1 led us to investigate the status of this histone mark in vivo. We found thatγH2A.X is detected in vivo in control embryos at E5.5 and more abundantly at E6.5, and is mainly localized to the nucleolus along with a diffuse nuclear pattern (Fig. 2C, D, S2A). Strikingly, Chd1 KO embryos entirely lack this γH2A.X signal at both developmental stages (Fig. 2E), while global H2A.X is retained (Fig. S2B). These findings in vivo are in agreement with the ES cell data above, showing they are not an artifact of cell culture. The early post-implantation epiblast is one of the fastest proliferating cell types in mammals, with doubling times between 2-8 hours^19^. Interestingly, the fastest rates of proliferation were recorded at E6.5, where we find high levels of nucleolar γH2A.X (Fig. 2 and S2) Overall, the results indicate that nucleolar accumulation of γH2A.X in vivo correlates with rRNA synthesis and proliferation rate, and all three of these are dependent on Chd1 (this study and ^3^).

### Double-stranded DNA breaks accumulate at rDNA in Chd1 KO ES cells

The loss of γH2A.X in Chd1 KO cells could simply be a consequence of the reduced global transcriptional output^3^ and therefore a lower occurrence of transcription-induced DNA breaks. We therefore set out to determine the levels and genomic location of DSBs in control vs Chd1 KO ES cells. We performed DSB labeling by terminal transferase followed by affinity purification and qPCR or deep sequencing (DSB-qPCR or DSB-seq)^17^ (Fig. 3A). We first focused on rDNA (Fig. 3B) due to the mainly nucleolar accumulation of several Chd1-interacting proteins (Figs. 1 and 2), as well as the reduced rRNA synthesis and nucleolar fragmentation observed in Chd1 KO cells^3^. Surprisingly, we found that deletion of Chd1 leads to an accumulation of DSBs at rDNA (Fig. 3C). DSBs are most abundant in the enhancer, promoter and 5’ end of the rDNA repeat, and decrease along the transcribed unit (Fig. 3C). We quantified nascent rRNA transcription by metabolic labeling of RNA with 5’-ethynyluridine (EU) coupled to biotin, followed by affinity capture and qPCR (EU-capture). Chd1 loss leads to a decrease in nascent rRNA transcripts, particularly at the 5’ end of the transcription unit (Fig. 3D). Thus, Chd1 KO ES cells accumulate DSBs at rDNA in the absence of any external DNA damage and in a context of reduced nascent rRNA transcription. Moreover, these results suggest that the loss of γH2A.X in Chd1 KO ES cells is not due to a lower level of DSBs, but rather to defective repair.

**Figure 3.**
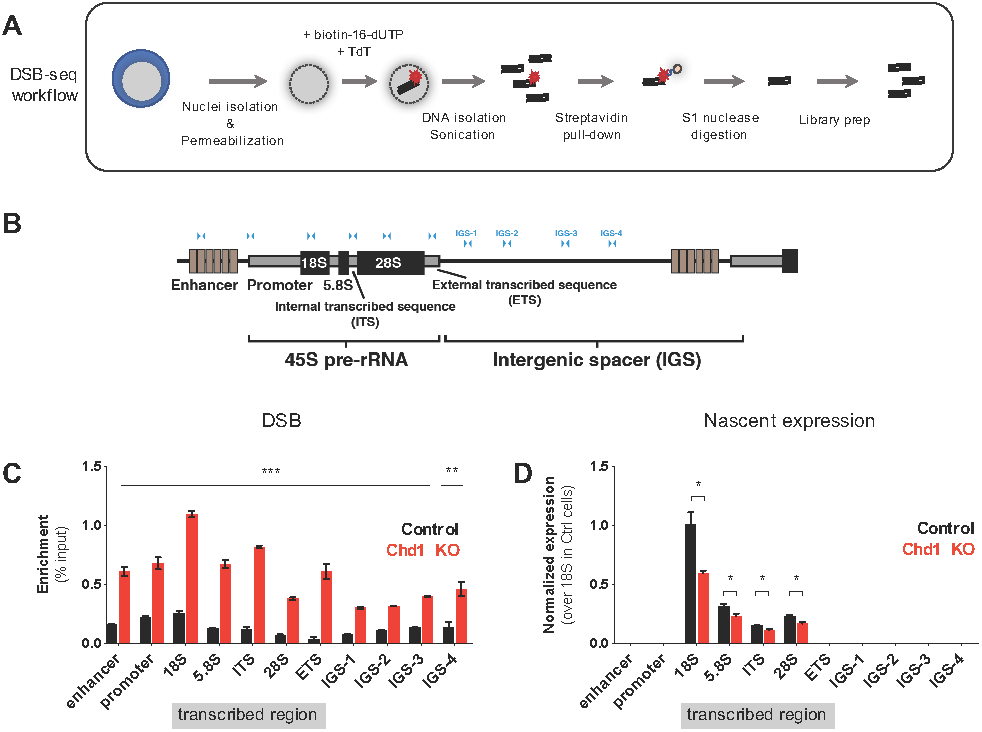
Double-stranded breaks accumulate at ribosomal DNA in Chd1 KO cells. A) Schematic depiction of the workflow of DSB-PCR/seq. B) Schematic depiction of one rDNA transcription unit. Primers are indicated with blue arrowheads. C) DSB levels along the rDNA unit in control and Chd1 KO cells, normalized to input. D) Nascent rRNA transcription in control and Chd1 KO cells. Error bars show standard deviation of two biological replicates. Statistical tests performed are T-tests with Holm-Sidak correction. *, **, *** correspond to p-values of <0.05, 0.01, 0.001.

### Chd1 loss induces widespread accumulation of double-stranded DNA breaks at RNA Pol II-transcribed genes

We previously reported that Chd1 deletion leads to global *hypo*transcription of both RNA Pol I and Pol II-transcribed genes^3^. The unexpected increase in DSBs at rDNA (Pol I-transcribed) in Chd1 KO ES cells (Fig. 3C) led us to explore the status of DSBs at Pol II-transcribed genes, using DSB-seq (see Methods for procedure). For comparative analyses, we also performed Chd1 and RNA Pol II ChIP-seq in Chd1-Flag knock-in ES cells^3^.

DSB-seq revealed a remarkable widespread accumulation of DSBs at promoter regions of RNA Pol II-transcribed genes in Chd1 KO cells, relative to controls (Fig. 4A-C). DSBs occur immediately downstream of the transcription start site (TSS), where Chd1 binding peaks in wild-type (WT) conditions (Fig. 4A-C). At promoter regions (+/-1kb of TSS), DSBs in Chd1 KO cells positively correlate with GC content (Spearman ρ= 0.80, *p* < 10^-300^), Chd1 (Spearman ρ = 0.71, *p* < 10^-300^) and RNA Pol II (Spearman ρ = 0.63, *p* < 10^-300^) binding in WT cells, and negatively correlate with nucleosome occupancy (Spearman ρ = -0.29, *p* < 5×10^-280^). The propensity to accumulate DSBs in Chd1 KO ES cells does not correlate with wild-type gene expression levels (Fig. 4B-D, Spearman ρ = 0.012, *p* > 0.306) or reduced expression upon Chd1 loss^3^ (Fig. S3A).

**Figure 4.**
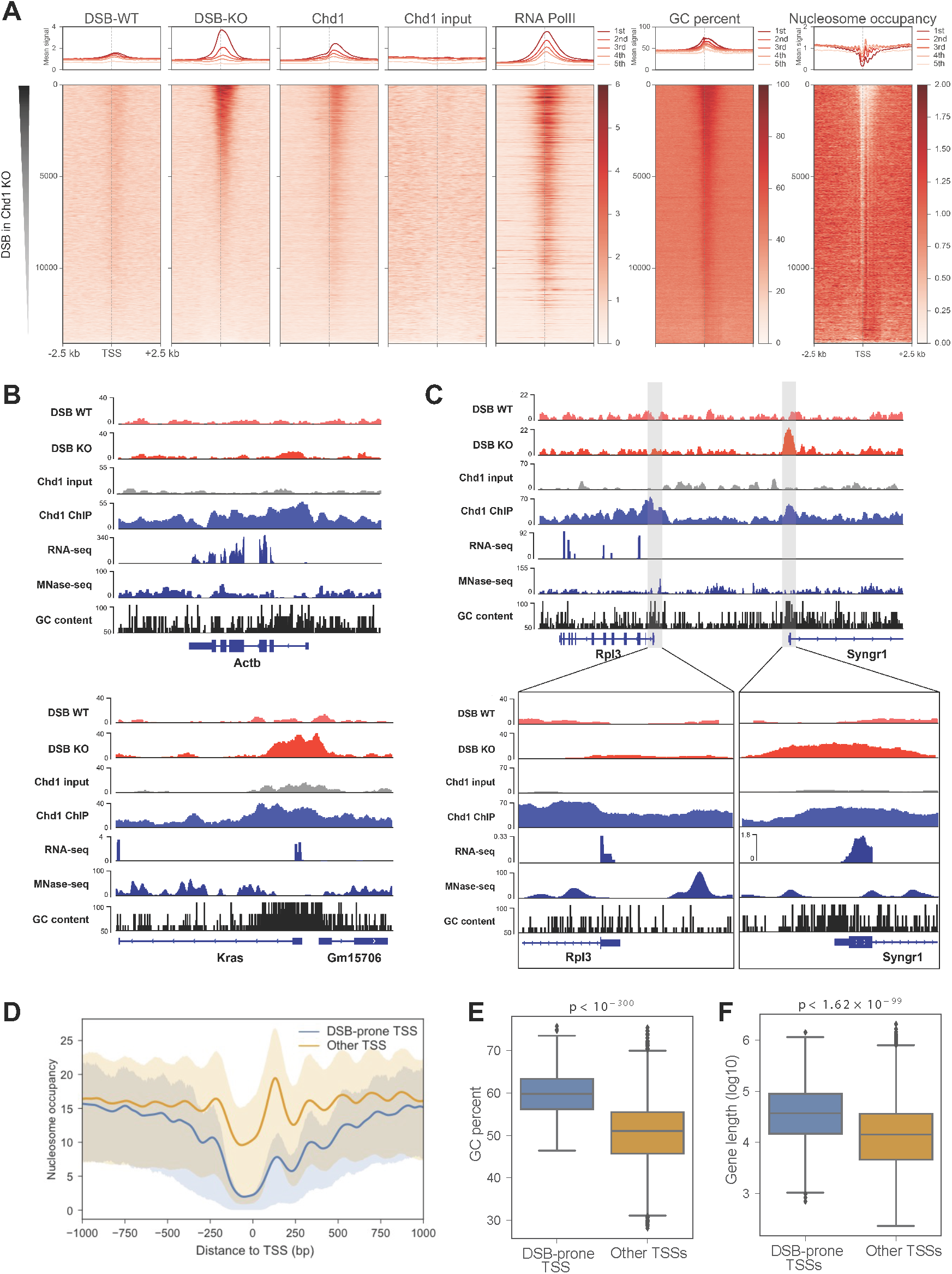
Widespread double-stranded break accumulation at TSSs in Chd1 KO cells. A) Heatmaps showing DSB, Chd1 binding, Pol II levels, nucleosome occupancy and GC content. See Methods for details. Upper panels show the mean signal within each quintile of genes sorted according to the same order as in the heatmaps. B, C) Genome browser views of DSB-prone (Kras, Syngr1) and other (Actb, Rpl3) genes. Note that DSB propensity does not correlate with expression level (See Fig. S3A). D) Nucleosomal patterns at TSSs of DSB-prone and non-DSB-prone genes. E, F) GC content and gene length of DSB-prone TSSs in comparison to non-DSB-prone TSSs. Statistical tests are Wilcoxon rank-sum tests. In D, E, and F, only protein-coding genes with unique TSSs are included.

We detected 5671 DSB peaks in Chd1 KO ES cells (vs. only 54 in control), among which 1825 peaks mapped around the TSSs (-1kb to +100bp) of 1785 genes (DSB-prone genes). DSB peaks are enriched at TSSs, 5’ untranslated regions and exons (Fig. S3B). GO analysis predicts that genes involved in transcription, chromatin modification and signaling are particularly prone to DSB in Chd1 KO ES cells (Fig. S3C). To understand why these genes might be especially susceptible to DNA breaks, we analyzed the chromatin structure at DSB-prone TSSs in comparison to non-DSB-prone TSSs. We utilized ES cell MNase-seq datasets generated by Voong et al. ^20^ (Fig. 4D). DSB-prone genes display a more open chromatin structure at TSSs, with less regular nucleosomes surrounding the start site. In particular, the +1 nucleosome is less abundant in DSB-prone genes (Fig. 4D). Taken into consideration that this graph depicts a population average of many cells going through the transcription cycle, the data suggest that the +1 nucleosome in DSB-prone genes is frequently displaced to expose naked DNA. Interestingly, Chd1 peaks at this +1 position (Fig. 4A), suggesting that it promotes eviction of this nucleosome, as previously reported in embryonic fibroblasts^10^, and in doing so facilitates DNA repair and transcriptional elongation.

To probe why these 1785 genes have a more exposed TSS region and are DSB-prone upon Chd1 loss, we investigated genomic features of these genes. DSB-prone genes have a significantly higher GC content in their promoter-proximal region compared to non-DSB-prone genes (Fig. 4A, E, Wilcoxon rank-sum test *p* < 10^-300^). Moreover, DSB-prone genes are on average longer than non-DSB-prone genes (Fig. 4F, Wilcoxon rank-sum test *p* < 1.62×10^-99^). Previous studies showed that longer genes are more DSB-prone in neurons, due to the increased Topoisomerase activity required to relieve the torsional stress that accumulates during DNA unwinding (reviewed in^16^). Taken together, our results suggest that Chd1 remodels nucleosomes at GC-rich promoters of long genes to facilitate DNA repair and repeated cycles of RNA Pol II elongation in hypertranscribing ES cells.

## Discussion

This work describes a novel role for the chromatin remodeler Chd1 in promoting transcription-mediated DNA break repair in pluripotent stem cells. Our results further point to a central role for rDNA at the nucleolus in the coordination between hypertranscription and DNA integrity in hypertranscribing ES cells. rRNA comprises ∼80% of the RNA being synthesized in ES cells and therefore represents both a major focal point of hypertranscription as well as a vulnerability to DNA breaks. We had previously implicated Kap1, generally thought to be a repressor, in rRNA transcription^21^, and this study further contributes to clarifying that role. The phosphorylated version of Kap1 may be key to the repair of transcription-associated DSBs, a possibility that deserves further exploration.

We propose that the accumulation of unrepaired DNA breaks in Chd1 KO cells compromises nascent transcriptional output and leads to the proliferation defects and ultimate developmental arrest of the rapidly expanding post-implantation epiblast. We note that the loss of Chd1 is not catastrophic to ES cells, despite the widespread occurrence of DSBs at GC-rich promoters of transcribed genes: Chd1 KO ES cells remain undifferentiated and capable of gene transcription, albeit with a lower transcriptional output and a self-renewal deficit. These results suggest that promoter DSBs are still eventually repaired in Chd1 KO ES cells, albeit at a lower rate, which compromises optimal nascent transcription and proliferation.

Repair of DSBs requires efficient signaling spearheaded by the Atm kinase, which we show is defective in Chd1 KO ES cells. Interestingly, Chd1-deficient human cancer cells have defective DNA repair when exposed to ionizing radiation or radiomimetic chemicals^22,23^. These cells show reduced H2A.X phosphorylation and are more sensitive to PARP inhibitors. The present study is the first report of Chd1-mediated DNA repair activity under native conditions without any external insult. The TSS sequence and chromatin landscape appear to be the main determinants of DSB generation in ES cells. DSBs were shown to be conducive to transcription in several cell types, in the absence of exogenous DNA damage^11-13,17^. Such studies point to a correlation between gene transcription and the occurrence of DNA breaks. Chd1 KO ES cells, on the other hand, have lower nascent transcription levels and higher incidence of DNA breaks. Thus, the DNA breaks we observe in Chd1 KO ES cells are not caused by increased transcription but rather by defective repair, a notion further supported by the loss of γH2A.X. We speculate that hypertranscribing, proliferating cells may have an increased dependence on Chd1 to balance high transcriptional output with DNA integrity. It will be of interest to explore the potential role of Chd1 in transcription-associated DNA break repair in other cell types, both in physiological stem/progenitor cells as well as in proliferating tumor cells. Of note, CHD1 is the second most frequently mutated gene in prostate cancer, after PTEN^24,25^. In agreement with our data, Dellino et al. recently showed that release of paused RNA Pol II in breast cancer cells induces DSBs preferentially at long genes and can lead to chromosomal translocations^26^.

In highly proliferative cells, replication stress can occur due to the collision of replication and transcription complexes, resulting in DNA breaks that are marked by γH2A.X^27^. It is unlikely that the high γH2A.X signal in wildtype ES cells and epiblast is strictly due to replicative stress because high γH2A.X is observed in all cells regardless of cell cycle stage. Moreover, Chd1 KO ES cells have higher incidence of DSBs and reduced γH2A.X (this study) without significant deviations in cell cycle stage proportions^3^. Taken together, our findings suggest that hypertranscription puts ES cells, and potentially other stem/progenitor cells, at risk of DNA breaks and genomic instability, but this is countered by Chd1-dependent DNA repair.

Why Atm signaling is defective in Chd1 KO ES cells is a central question that remains elusive. In Chd1 KO cancer cells, γH2A.X is reduced due to reduced incorporation or retention of H2A.X at damage sites^23^. We did not observe global or local H2A.X loss in Chd1 KO ES cells or embryos. In contrast, both Atm and H2A.X levels are increased at rDNA upon loss of Chd1. Thus, the enzyme and its substrate are elevated at chromatin in Chd1 KO ES cells, but H2A.X phosphorylation is either defective or not sustained. It is possible that the chromatin remodeling activity of Chd1 facilitates access of Atm to its target site on H2A.X, or that other interactors of Chd1 promote Atm activity towards H2A.X. For example, Chd1 also interacts with PARP1 and the histone acetyltransferase Tip60 (data not shown), and both histone ADP ribosylation and acetylation are involved in chromatin relaxation at DNA repair sites ^28,29^. Further biochemical studies of how the activity of Atm and other aspects of DNA repair are modulated by Chd1 will shed light on the mechanisms by which stem and progenitor cells can undergo hypertranscription while preserving DNA integrity.

## ACKNOWLEDGMENTS

We thank members of the Santos Lab for input and critical reading of the manuscript, Richard Lao and members of the UCSF Institute of Human Genetics for assistance with sequencing and sonication. Samples were sequenced at UCSF Institute of Human Genetics Core Facility and Center for Advanced Technology, which is supported by the NIH (5P30CA082103). This research was supported by grants from Bloodwise and Medical Research Council plus the CRUK Manchester Centre award (C5759/A25254) to ADW who is supported by the NIHR Manchester Biomedical Research Centre, 2R01CA163336 to J.S.S and by NIH R01GM113014, R01GM123556 and Canada 150 Research Chair in Developmental Epigenetics to M.R.-S.

## AUTHOR CONTRIBUTIONS

A.B.-K. and M.R.-S. conceived of the project. A.B.-K. designed and performed the majority of experiments, with the following exceptions: M.G.-A. isolated, stained and imaged the embryos, A.J.K.W performed the mass spectrometry experiments, H.J. and M.H. performed bioinformatics analyses, under the supervision of J.S.S. A.D.W supervised the mass spectrometry experiments. M.R.-S. supervised the project. A.B.-K., and M.R.-S. wrote the manuscript with feedback from all authors.

## ACCESSION NUMBERS

Data have been deposited in Gene Expression Omnibus (GEO) under accession number GSE132137.

## EXPERIMENTAL PROCEDURES

### Mice

Chd1^Δ/+^ females (6-to 12-week-old) and males (6 week-to 6-month-old) were used to recover the embryos. Animals were maintained on 12 h light/dark cycle and provided with food and water *ad libitum* in individually ventilated units (Techniplast at TCP, Lab Products at UCSF) in the specific-pathogen free facilities at UCSF. All procedures involving animals were performed in compliance with the protocol approved by the IACUC at UCSF, as part of an AAALAC-accredited care and use program (protocol AN091331-03). Chd1 heterozygous mice were mated, embryos were collected at embryonic day 5 (E5.5) or day 6 (E6.5) after detection of the copulatory plug by dissecting uteri of pregnant females as described before^3^.

### ES cell culture

Chd1-Flag knock-in, Chd1^fl/Δ^ and Chd1^Δ/Δ^ ES cells were used as described^3^. Cells were grown in DMEM GlutaMAX with 15% FBS (Atlanta Biologicals), 0.1 mM non-essential amino acids, 50 U/ml Penicillin/Streptomycin (UCSF Cell Culture Facility), 0.1 mM EmbryoMax 2-Mercaptoethanol (Millipore) and 2000 U/ml ESGRO supplement (LIF, Millipore or Gemini) under ambient air with 5% CO_2_. Cells tested negative for mycoplasma contamination.

### Immunoprecipitation

Wild type E14 or Chd1-Flag knock-in cells were used. Cells were fractionated into cytoplasmic and nuclear compartments prior to immunoprecipitation. For cytoplasmic extracts, cells were lysed in Buffer A (10mM Hepes pH7.9, 5mM MgCl_2_, 0.25M Sucrose and 0.1% NP-40 supplemented with 1x Halt protease inhibitor cocktail (Thermo Fisher Scientific, 78425), 1 mM PMSF, 5 mM NaF and 1 mM NaVO_4_) and were centrifuged for 10 min, 4400 rpm at 4°C. For nuclear extracts, the resulting pellets were resuspended in buffer B (10mM Hepes pH 7.9, 1mM MgCl2, 0.1mM EDTA, 25% glycerol, 0.5% Triton X-100 and 0.5M NaCl supplemented with 1x Halt protease inhibitor cocktail (Thermo Fisher Scientific, 78425), 1 mM PMSF, 5 mM NaF and 1 mM NaVO_4_) and homogenized by passing through 18Gx1 1/2’’ size needles. Nuclear extracts were quantified using Pierce BCA Protein Assay kit (23225). For immunoprecipitation, 100 μg of extract was adjusted to 500 μl total volume and 150 mM final NaCl concentration, and incubated in the presence of 20 μl pre-washed Protein A or G Dynabeads (Thermo Fisher Scientific, 1002D or 1004D) and the following antibodies: Flag (Sigma, F1804), Kap1 (Abcam, ab22553), Atm phospho S1981 (Active Motif, 39529), Atr (Santa Cruz Biotechnology, sc-1887), Parp1 (Santa Cruz Biotechnology, sc-25780), Top2β (Santa Cruz Biotechnology, sc-13059), Xrcc1 (Santa Cruz Biotechnology, sc-11429). Beads were washed three times in buffer B, then boiled in 2x Laemmli Buffer with 5% β-mercaptoethanol. Western blot was performed as described below.

### IP-Mass Spectrometry

Chd1-Flag ES cells were lysed and processed as above. Nuclear extracts were used for immunoprecipitation. For mass spectrometry, Flag IP and beads control were run on a denaturing gel. Gel was stained using Coomassie Brilliant Blue. IP and control lanes were cut into 10 pieces avoiding light and heavy antibody chains and frozen until processed for mass spectrometry.

Protein bands were excised, destained with repeated incubation in 200 mM ammonium bicarbonate, 40% [v/v] acetonitrile. Gel pieces were dried with three washes in 100% acetonitrile and then trypsinised (Trypsin resuspended in 100 mM ammonium bicarbonate, 5% [v/v] acetonitrile) overnight at 37°C. Peptides were extracted from the gel pieces by incubation in 50% [v/v] acetonitrile, 0.1% [v/v] formic acid, peptides were desiccated and resuspended in 2% [v/v] acetonitrile, 0.05% [v/v] trifluoroacetic acid; pH 2.7. For each analysis, 10% of the peptide sample was loaded onto an Acclaim Pepmap C18 Trap (500 µm x 5 mm) and flow was set to 30 µl/min of 2% [v/v] acetonitrile, 0.05% [v/v] trifluoroacetic acid for 5 min. Analytical separation of the peptides was performed using Acclaim PepMap100C18 Column (3 µm, 75 µm x 500 mm) on a U3000 RSLC (Thermo). Briefly, peptides were separated over a 91 minutes solvent gradient from 2% [v/v] acetonitrile, 0.1% [v/v] formic acid to 40% [v/v] acetonitrile, 0.1% [v/v] formic acid on-line to a LTQ Orbitrap Velos (Thermo). Data was acquired using an data dependant acquisiton (DDA) method where, for each cycle one full MS scan of m/z 300 - 1700 was acquired in the Orbitrap at a resolution of 60,000 at m/z 400 with an AGC target of 1×10e6. Each full scan was followed by the selection of the 20 most intense ions, CID and MS/MS analysis was performed in the LTQ. Selected ions were excluded from further analysis for 60 seconds. Ions with an unassigned charge or a charge of +1 were rejected.

Data were analysed using Mascot (Matrix Sciences) the parameters were; Uniprot database, taxonomy *Mus Musculus*, trypsin with up to 1 missed cleavage allowed, variable modification were oxidised methionine, phosphorylated serine, threonine and tyrosine and the peptide tolerance of 0.025 Da and 0.03 Da for MS/MS tolerance.

### Western blot analysis

Chd1^fl/Δ^ and Chd1^Δ/Δ^ (7 days post-induction of KO) ES cells were used. For analysis of whole cell extracts, cells were lysed in RIPA buffer (150 mM NaCl, 1% NP-40, 0.5% sodium deoxycholate, 0.1% SDS, 50 mM Tris pH 8.0, 1x Halt protease inhibitor cocktail (Thermo Fisher Scientific), 1 mM PMSF, 5 mM NaF and 1 mM NaVO_4_). Cells were incubated for 30 minutes on ice, then sonicated using a Bioruptor (Diagenode) for 5 minutes with settings high, 30 seconds on, 30 seconds off. Laemmli Buffer with 5% β-mercaptoethanol was added to 1x and samples were boiled at 95C for 5 min. Extracts were loaded into 4-15% Mini-Protean TGX SDS Page gels (Bio-Rad). Proteins were transferred to PVDF membranes. Membranes were blocked in 5% milk/PBS-T buffer for 30 min and incubated either overnight at 4°C or for 1 hour at room temperature with the following antibodies: Chd1 (Cell Signaling, 4351), Top2β (Santa Cruz Biotechnology, sc-13059), Atm (Genetex, GTX70103), Kap1 (Abcam, ab22553), Nucleolin (Abcam, ab22758), Polr1a (Cell Signaling, D6S6S), H2A.X (Abcam, ab11175), γH2A.X (Abcam, ab2893), Gapdh (Millipore, MAB-374), anti-rabbit/mouse/goat secondary antibodies (Jackson Labs, 115-035-062, 111-035-144). Membranes were incubated with ECL or ECL Plus reagents and exposed to X-ray films (Thermo Fisher Scientific).

### Immunofluorescent staining and imaging

Chd1^fl/Δ^ and Chd1^Δ/Δ^ ES cells were used. Cells were plated on matrigel in 8-chamber polystyrene vessels. Cells were fixed in 4% paraformaldehyde for 10 minutes, washed with PBS and permeabilized with 0.2% Triton X-100 in PBS for 5 minutes on ice. After blocking in PBS, 2.5% BSA, 5% donkey serum for 1 hour, cells were incubated overnight at 4°C with the following antibodies: Kap1 (Abcam, ab22553), phospho S824 Kap1 (Abcam, ab70369), γH2A.X (Abcam, ab2893), Nucleolin (Abcam, ab22758), Atm (Genetex, GTX70103), Top2β (Santa Cruz Biotechnology, sc-13059). Cells were washed in PBS-Tween20, 2.5% BSA, incubated with fluorescence-conjugated secondary antibody (Life Technologies) for 2 hours at room temperature and mounted in VectaShield mounting medium with DAPI (Vector Laboratories). Imaging was performed using a Leica BL-23 microscope. Staining and imaging of E5.5 and E6.5 embryos were performed as described before^3^. A minimum of 3 embryos was used for each experiment.

### Chromatin Immunoprecipitation (ChIP)

Chd1^fl/Δ^ and Chd1^Δ/Δ^ (7 days post-induction of KO) ES cells were used. ChIP was performed as described before ^30^. After aspiration of culture medium, cells were washed with PBS and fixed on the culture dish using 1% formaldehyde in PBS for 10 minutes at room temperature (RT). Glycine was added to a final concentration of 125 mM to quench formaldehyde for 5 minutes at RT. Cells were washed twice with ice-cold PBS, incubated in Swelling Buffer (25 mM HEPES pH 7.9, 1.5 mM MgCl2, 10 mM KCl, 0.1% NP-40 with 1x Halt protease inhibitor cocktail (Thermo Fisher Scientific, 78425), 1 mM PMSF, 5 mM NaF and 1 mM NaVO_4_) for 10 minutes, scraped, passed through an 18Gx11/2” needle (5x) and spun down at 3,000g, 4°C, 5 minutes. Nuclei were resuspended in Sonication Buffer (50 mM HEPES pH 7.9, 140 mM NaCl, 1mM EDTA, 1% Triton X-100, 0.1% Na-deoxycholate 0.1% SDS with 1x Halt protease inhibitor cocktail, 1 mM PMSF, 5 mM NaF and 1 mM NaVO_4_) and sonicated using a Covaris S2 sonicator with settings 5% duty cycle, intensity 4, cycles per burst 200, frequency sweeping. 20 μl chromatin was incubated sequentially with 1 μl RNaseA and 5 μl proteinase K in 100 μl total volume at 37°C for 30 min and 65°C for 1h, purified using a Qiagen PCR purification kit and DNA content was quantified using a NanoDrop. Fragment size distribution was checked on a 1% agarose gel. Chromatin was snap frozen if not immediately used for IP. Chromatin volume equivalent to 25 μg DNA was used for each IP. Chromatin was immunoprecipitated in the presence of 20 μl pre-washed Protein A or G Dynabeads (Thermo Fisher Scientific, 1002D or 1004D) and the following antibodies: Flag (Sigma, F1804), Kap1 (Abcam, ab22553), phospho S824 Kap1 (Abcam, ab70369), γH2A.X (Abcam, ab2893), H2A.X (Abcam, ab11175), H1 (Thermo Fisher Scientific, PA128374), Polr1a (Cell Signaling, D6S6S), Atm (Genetex, GTX10701). Beads were washed in sonication buffer (2 times), wash buffer A (sonication buffer with 500 mM NaCl) and TE buffer (10 mM Tris pH 8.0, 1 mM EDTA), and resuspended in elution buffer (50 mM Tris pH 7.5, 1 mM EDTA, 1% SDS) with 1 μl RNaseA and 5 μl proteinase K in 100 μl total volume. After incubation at 37°C for 30 min and 65°C for 2h to overnight, DNA was purified using a Qiagen PCR purification kit. qPCR was performed with KAPA SYBR FAST qPCR Master Mix (Kapa Biosystems) and amplified on a 7900HT Real-time PCR machine (Applied Biosystems). For sequencing, libraries were prepared using the NEBNext ChIP-seq Library Prep for Illumina kit. Library quality and quantity were analyzed using Bioanalyzer (Agilent). Samples were sequenced on a HiSeq 2500 using single-end 50 bp sequencing reads in rapid mode.

### Detection of double-stranded DNA breaks

Chd1^fl/Δ^ and Chd1^Δ/Δ^ ES cells were used. For detection of double-stranded DNA breaks, we combined the Baranello et al^17^ and BLESS protocols^31^ to achieve end labeling in situ. 1.5×10^8^ cells were fixed in 2% formaldehyde suspension for 30 min at room temperature, followed by quenching in 0.25M glycine for 5 min. Cells were centrifuged, washed twice in PBS, and incubated in 25 ml lysis buffer (10 mM Tris pH 8.0, 10 mM NaCl, 1 mM EDTA, 1 mM EGTA, 0.2% NP-40) for 1 h at 4°C. Pellets were resuspended in nucleus break buffer ((10 mM Tris pH 8.0, 150 mM NaCl, 1 mM EDTA, 1 mM EGTA, 0.3% SDS, 1mM DTT) and incubated for 45 min at 37C. After centrifugation, nuclei were resuspended in 1x NEB buffer 2 on ice. 10 μl Proteinase K (20 mg/ml) was added, digestion was performed for 4 min at 37C, and samples were immediately returned to ice and 25 μl PMSF was added. Nuclei were centrifuged, and resuspended in 5 ml 1x NEB Buffer 2 + 15 μl Triton X-100, and centrifuged again at 200g for 10 min at 4°C. Nuclei were washed once in water, and divided into two tubes for end labeling and control reactions. Nuclei were resuspended in 625 μl 1x TdT buffer with 2.5 μl TdT (Promega, M1871) and 2 μl biotin-16-dUTP (Roche 11093070910), and were incubated at 37°C for 1 hour. Control reaction was assembled the same way without TdT. 1:50 volume of 0.5M EDTA was added to stop the reaction. Proteins were digested with 5 μl Proteinase K (20 mg/ml) at 2h to overnight at 65°C. Labeled DNA was precipitated using sodium acetate (70 μl) and isopropanol (700 μl), centrifuged, washed with 70% ethanol and resuspended in water. Genomic DNA was sonicated using a Covaris S2 sonicator with settings 5% duty cycle, intensity 4, cycles per burst 200, frequency sweeping for 4.5 cycles. DNA amount was quantified on a NanoDrop. Size distribution was checked on a 1% agarose gel. For biotinylated DNA pull-down, 20 μg DNA was adjusted to 600 μl final volume in W&B buffer (10 mM Tris pH 7.5, 1 mM EDTA, 1 M NaCl). 10 μl Dynabeads MyOne C1 was washed twice with W&B buffer and added to DNA. Sample was incubated for 1 hour at 4°C. Beads were washed twice in W&B buffer, and resuspended in 100 μl elution buffer (95% v/v formamide, 10 mM EDTA). After incubation at 65°C for 5 min, DNA was purified using a Qiagen PCR purification column. For sequencing, biotinylated overhangs were removed using S1 nuclease. Sequencing libraries were prepared using the NEBNext ChIP-seq Library Prep for Illumina kit. Library quality and quantity were analyzed using Bioanalyzer (Agilent). Samples were sequenced on a HiSeq 4000 using single-end 50 bp sequencing reads.

### Nascent RNA capture followed by qRT-PCR

To measure nascent transcriptional changes at specific loci, ES cells were analyzed using the Click-iT Nascent RNA Capture Kit (Life Technologies). Chd1^fl/Δ^ and Chd1^Δ/Δ^ cells were incubated with 0.2 μM 5-ethynyl uridine (EU) for 30 minutes to label nascent transcripts. Cells were washed, harvested by trypsinization and counted. Total RNA was isolated from the 10^6^ Chd1^fl/Δ^ or Chd1^Δ/Δ^ cells using the Qiagen RNeasy Mini Kit (Qiagen) and processed according to manufacturer’s instructions. qPCR was performed with KAPA SYBR FAST qPCR Master Mix (Kapa Biosystems) and amplified on a 7900HT Real-time PCR machine (Applied Biosystems).

### Bioinformatic analyses

#### ChIP-seq, DSB-seq, and MNase-seq data processing

ChIP-seq and DSB-seq reads were mapped to the mm9 genome using Bowtie2^32^ with the options --end-to-end --sensitive --score-min L,-1.5,-0.3. Reads with mapping quality smaller than 13 were filtered out. PCR duplicates were removed by keeping at most one mapped read at each genomic position. Biological replicates were then combined. Read coverage profiles were generated using bedtools^33^ after extending the mapped reads from 5’ to 3’ end to 200 bp. Processed MNase-seq data in mouse ES cells were obtained from Voong et al. ^20^ with accession number GSE82127. Center-weighted nucleosome occupancy was calculated from the provided nucleosome center scores as described in Voong et al. and used throughout the paper.

#### DSB-seq peak calling and annotation

DSB-seq peaks were called using MACS2^34^ with the options --shift 0 --nomodel --extsize 200 -g mm and default q-value cutoff 0.05. 5903 and 183 peaks were identified for Chd1 KO and WT cells, respectively. After filtering out peaks overlapping with blacklisted regions (http://mitra.stanford.edu/kundaje/akundaje/release/blacklists/mm9-mouse/mm9-blacklist.bed.gz), 5671 and 54 peaks were retained for Chd1 KO and WT cells, respectively. The identified peaks were associated with genomic annotations using HOMER annotatePeaks.pl^35^. In particular, 1825 peaks in Chd1 KO cells were mapped around TSSs (-1kb to +100bp) of 1785 genes (DSB-prone genes). In Fig 4A, genes were ranked based on mean DSB levels in Chd1 KO cells within +/-1kb of TSS. Values in each heatmap are divided by the mean of the entire matrix (except for GC content), respectively, and then smoothed using a Gaussian kernel (5 genes by 20 bp). Only protein-coding genes with unique TSSs were included to avoid ambiguity. The identified peaks were associated with genomic annotations <u>(HOMER mm9.v5.7)</u> using HOMER annotatePeaks.pl.

#### Gene annotations

RefSeq gene annotations were downloaded from UCSC Table Browser. In order to avoid ambiguity, only protein-coding genes with unique TSSs in autosomes were used. Genes whose TSSs +/-10kb overlap with blacklisted regions were further filtered out. As a result, 14109 genes were retained and used throughout the paper. In Figure 4B, we further required that the gene has a unique expression value provided by Guzman-Ayala et al. ^3^ (GSE57609), resulting in 7524 genes. In Figure 4D-F, 1107 DSB prone genes (intersection between the 1785 genes identified by HOMER and the 14109 filtered genes) were compared with 13002 (=14109-1107) non-DSB-prone genes. In Figure 4F, the mean gene length of different isoforms was used for genes with different transcription termination sites (TTS). Gene ontology analysis was performed using the DAVID software^36,37^.

## SUPPLEMENTARY FIGURES

**Figure S1.**
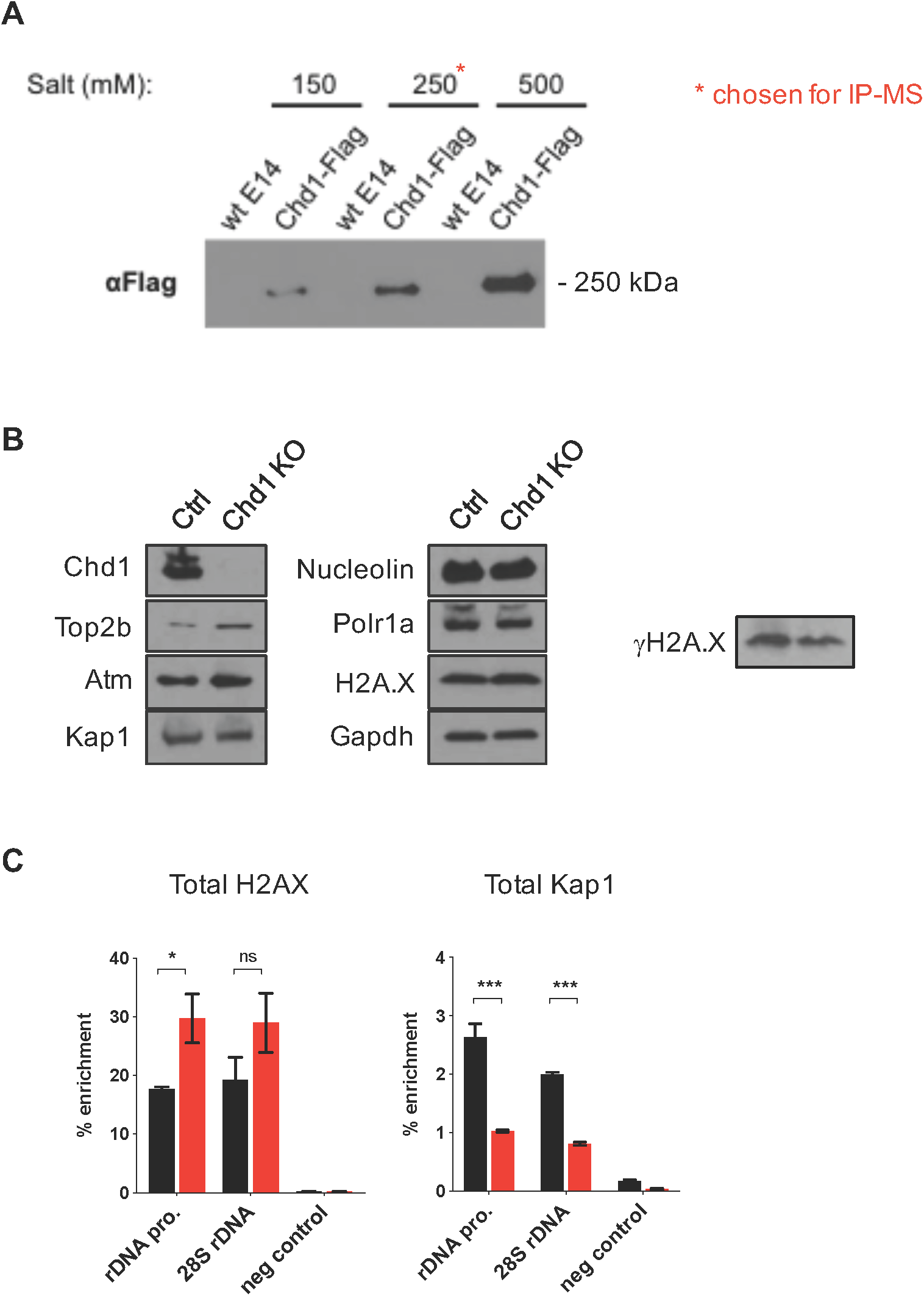
Further characterization of DNA repair protein abundance in Chd1 KO cells. A) Expression levels of indicated proteins in control and Chd1 KO ES cells. Whole cell extracts were used. B) Quantification of total H2A.X and Kap1 levels at rDNA in control and Chd1 KO cells by ChIP-PCR.

**Figure S2.**
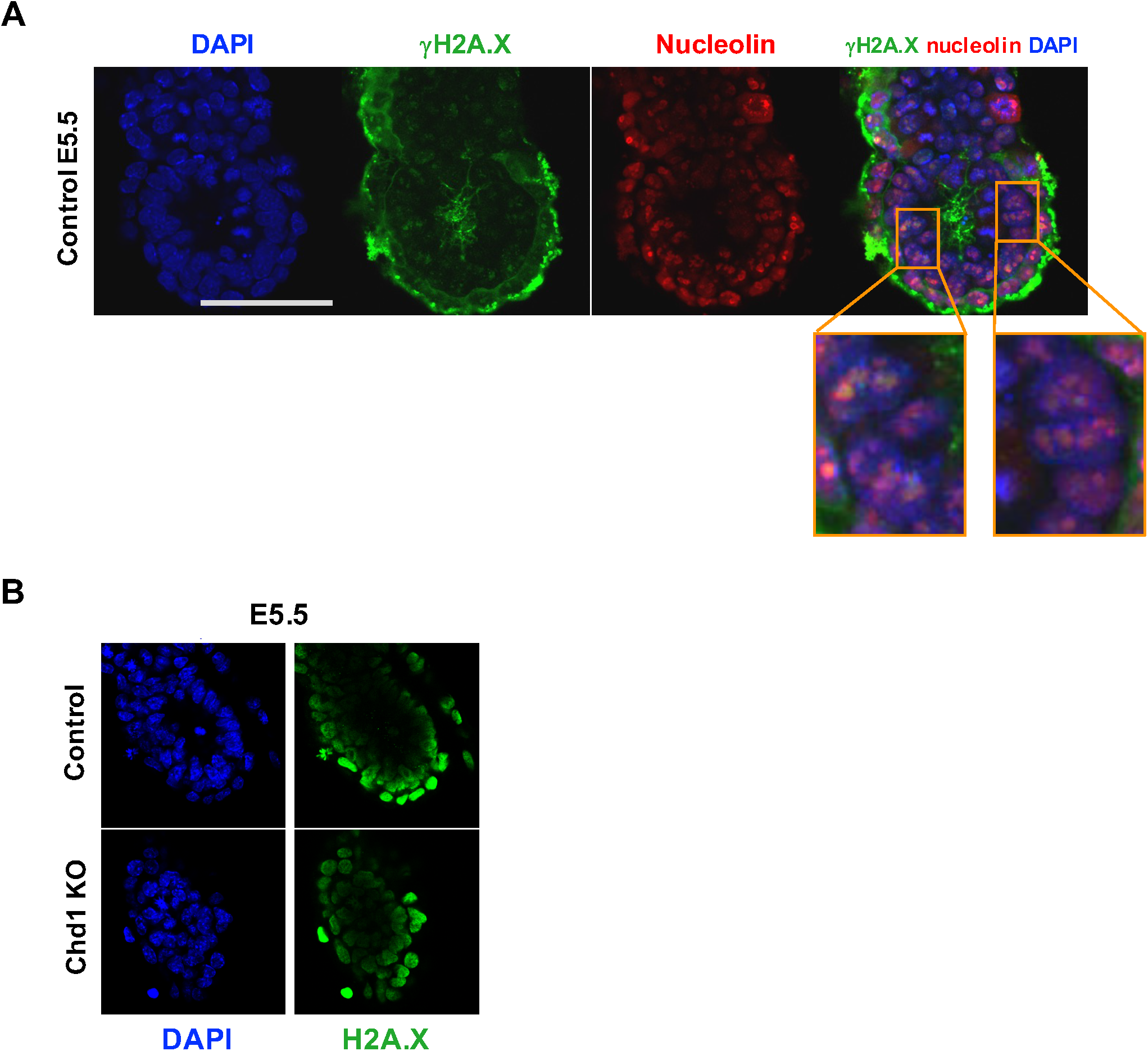
Immunofluorescent staining of A) γH2A.X and Nucleolin B) total H2A.X in Chd1 KO and control embryos at E5.5. Total H2A.X remains unchanged upon Chd1 deletion. γH2A.X signal colocalizes with Nucleolin.

**Figure S3.**
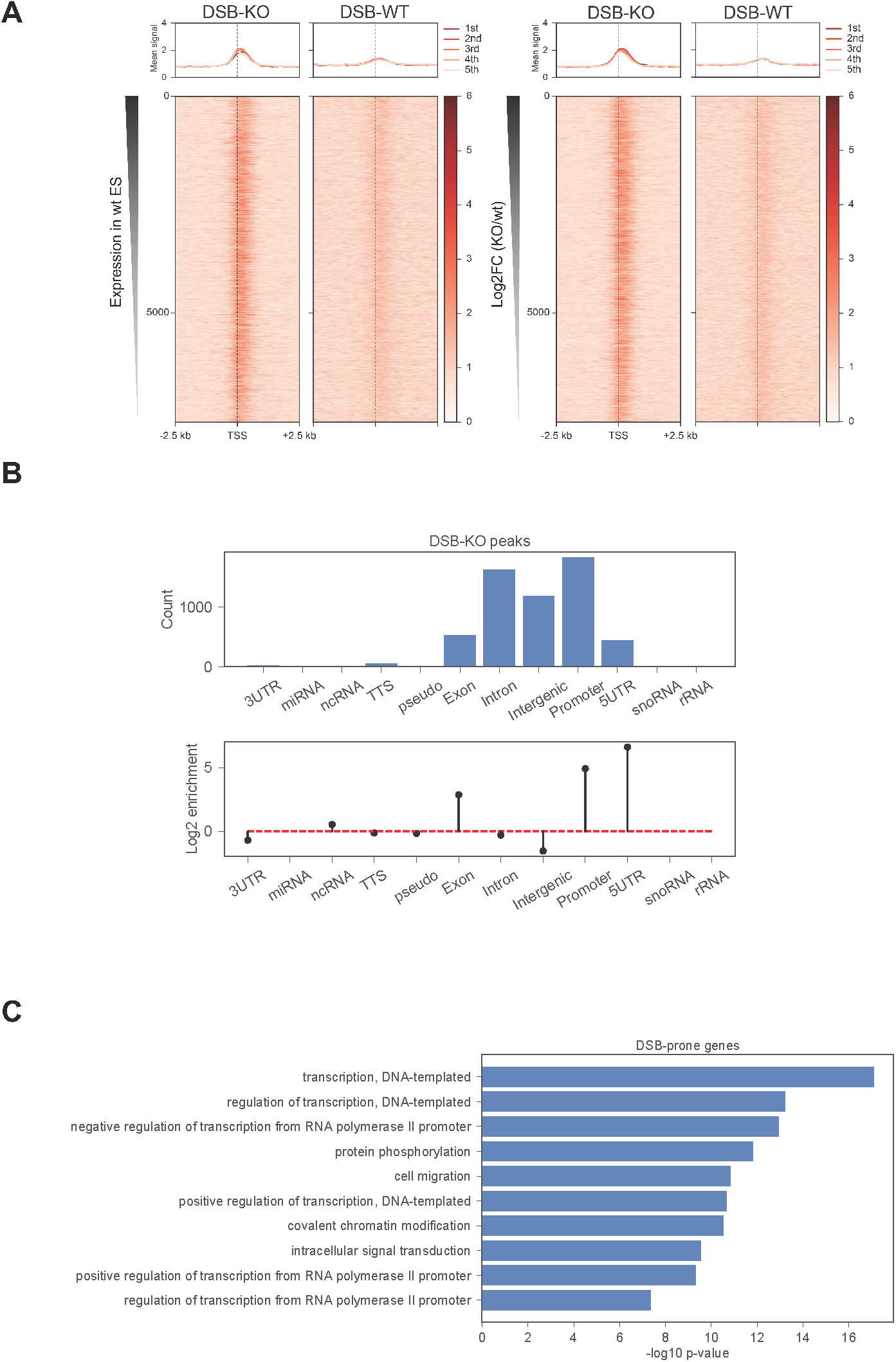
Characterization of DSB-prone genomic regions and associated gene functions. A) Heatmaps showing DSB levels in wt and Chd1 KO cells. Genes are ranked based on expression levels in wt cells (left panel) or fold change in Chd1 KO vs. wt cells (right panel). Only protein-coding genes with unique TSSs and provided expression values from Guzman-Ayala et al. ^3^ are included. B) Distribution and enrichment of DSBs on various genomic annotations. The log2 enrichment was calculated as the log2 ratio of the fraction of peaks associated with a specific genomic annotation and the fraction of the genome assigned to the same genomic annotation, as returned by HOMER. C) Gene ontology pathways associated with 1785 DSB-prone genes.

